# Channel opening duration in adult muscle nAChRs determined by activated external ACh binding site

**DOI:** 10.1101/2021.11.15.468683

**Authors:** Dmitrij Ljaschenko, Achmed Mrestani, Martin Pauli, Josef Dudel, Manfred Heckmann

**Author notes:** Equal contributions (Dmitrij Ljaschenko, Achmed Mrestani, Martin Pauli).

## Abstract

We recorded currents through the cell membrane at single nAChR molecules, held at ACh or Epibatidine (Ebd) concentrations of 0.01, 0.1, 1, 10 or 100 µM. The measured current amplitudes had an absolutely fixed value of 15 pA. This was valid for different agonists at all concentrations. Binding an agonist at one or both sites in the ring of subunits allowed to open the channel, the site that initiated the opening determined the duration of the final opening of the channel. In addition, the current flow was continuously interrupted by < 3 µs shut times. The resolution of our records was optimized to reach 5 µs, but was insufficient to resolve an unknown proportion of shorter shut times. Therefore, measured durations of openings are overestimated, and cited in brackets: τ_o1_ (3 µs) elicited by agonist-binding at the δ-site, τ_o2_ and τ_o3_ (40 and 183 µs) by binding at the ε-site, and τ_o4_ (752 µs) by binding at the ε- and δ-site. Mono-liganded nAChRs trigger short bursts of 0.6 ms duration. Bi-liganded nAChRs generate long bursts that at low agonist concentrations last 12 ms. Above 10 µM ACh, long bursts are shortened, with 100 µM ACh, to 5 ms, and further at higher concentrations. While ACh was the main agonist, Ebd bound more effectively than ACh to the ε-site.

**SIGNIFICANCE:** Transmembrane currents were recorded from single nAChR molecules. Binding ACh at one of the external receptor sites opens the central current channel repeatedly for a period selected by the activated receptor site, and transmembrane current can flow with fixed 15 pA amplitude (same at various agonists and range of concentrations, unaffected by desensitization). Current flow is interrupted by fixed < 3 µs blocks of the central channel that border each opening. Sets of current block and opening can be repeated for receptor-determined periods of time and form bursts of channel openings. The results may help to interpret the structural reactions in the center of the molecule that are addressed in the introduction.

## INTRODUCTION

Muscle-type nicotinic acetylcholine (ACh) receptor channels (nAChRs) are members of the pentameric ligand gated ion channel family. Receptors of this family are made up of five identical or homologous symmetrically arranged subunits filling a cylinder at least 12 nm in height and 7 nm in diameter whose rotational axis coincides with the central ion pathway (Nemecz et al., 2016, Rahman et al., 2020). Muscle-type nAChRs have a precise stoichiometry with two α-, a single β-, a single δ- and a single ε-subunit in adult receptors. The two α-subunits of muscle-type nAChRs form two distinct ligand binding sites at interfaces with adjacent δ- or ε-subunits. The binding pocket is lined by several highly conserved aromatic residues. Here ligand induced conformational changes occur which are transmitted from the extracellular domains to the transmembrane domains (TMDs), resulting in the opening of the channel. In a combination of equilibrium and nonequilibrium molecular dynamics simulations of human α4β2 receptors these conformational changes proceed, firstly through loop C in the principal subunit, and are subsequently transmitted, gradually and cumulatively, to loop F of the complementary subunit, and then to the TMDs through the M2-M3 linker (Oliveira et al., 2019).

Since early kinetic studies of single nAChRs closed times around the dead time have been important in the quest to decipher receptor activation (Colquhoun & Sakmann 1985; Sine & Steinbach, 1987; Auerbach & Lingle, 1987). More recently, briefest closed times, in terms of mechanism, were suggested to originate from a preopen closed state, known as “flipped” or “primed”, in the path toward channel opening (Lape et al., 2008; Lape et al., 2009; Mukhtasimova et al., 2009; Mukhtasimova et al., 2016).

Muscle-type nAChRs generate, in addition to bursts of bi-liganded openings, short mono-liganded openings (for a recent review see Bouzat & Sine, 2018). Furthermore, δ- and ε-sites contribute unequally to gating of muscle-type nAChRs (Akk *et al*., 1996; Nayak *et al*., 2014; Nayak *et al*., 2016; Mukhtasimova et al., 2016).

To our knowledge, the precise functions of the two binding sites for ACh on the surface of adult muscle fibers have not yet been identified. This task will be the first aim of the present study. Binding of ACh to one of the sites may initiate the opening of the transmembrane channel of the nAChR molecule. Then we shall investigate the relations between surface receptor and channel opening. The reactions within the molecule proceed at rates beyond our temporal resolution. It seems impossible to derive a reaction scheme presently, but we propose a simple network of states of the molecule that may describe the functions.

## MATERIALS AND METHODS

### Receptor expression in cell culture

HEK 293 cells were cultured on poly-L-lysine coated coverslips at 37 °C, 5 % CO_2_, in Dulbecco’s modified Eagle’s medium (Gibco). The medium was supplemented with 10 % fetal calf serum (Gibco), 100 units/ml penicillin and 100 µg/ml streptomycin (Gibco, Heckmann *et al*., 1996). To express murine adult nAChRs, cells were transiently transfected employing the calcium phosphate co-precipitation method (Groot–Kormelink *et al*., 2002). The mouse muscle acetylcholine receptor subunit ε was encoded on a pcDNA3 plasmid (Akk, 2002), the subunits α, β, δ on three pRc/CMV plasmids (ratio α, β, δ, ε: 2, 1, 1, 1). In order to mark transfected cells, the GFP carrying plasmid pmaxGFP (Amaxa) was added to the transfection solution.

### On-cell patch clamping

Prior to recordings, coverslips were transferred to recording chambers, which contained a physiological solution (in mM, 162 NaCl, 5.3 KCl, 2 CaCl_2_, 0.67 NaH_2_PO_4_, 15 HEPES, pH 7.4). Patch pipettes contained additionally either acetylcholine (ACh) at concentrations of 0.01, 0.1, 1, 10 and 100 µM, or epibatidine (Ebd) at 0.01, 0.1 and 1 µM. On-cell patch clamp recordings were performed at room temperature, (19.5–22.9 °C, mean: 21.7 ± 1.0 SD °C). To improve the signal to noise ratio all patches were voltage clamped at -200 mV, while the resting potential of transfected HEK cells was around -30 mV.

In some experiments δ-sites were blocked. To this end, cells were incubated in 1 µM *α-Conotoxin M1* (CTx; Sine *et al*., 1995; Cortez *et al*., 2007; Azam & McIntosh, 2009; Stock *et al*., 2014) containing culture medium for 10 minutes at 37 °C. After transferring the cover slip to the recording chamber, a CTx free pipette was used for recordings. Since 50 - 150 mbar pressure was applied to each pipette prior to patch formation, access of CTx from bath solution to the patch was prevented. In most patches all δ-sites were blocked and recording could start. This procedure was usefull, since the concentration of CTx had to be high enough to block all δ-sites rapidly, but further application of CTx to the patch would have blocked too many ε-sites. During the recording CTx dissociated from the δ-site at a very low rate. Eventually agonists could bind to both binding sites, eliciting long bursts. Thus, as soon as the first long burst appeared, evaluation of the recording was stopped.

To block the ε-site of the receptor 1 µM Waglerin-1 (WTx) was applied (McArdle *et al*., 1999, Molles *et al*., 2002, Teichert *et al*., 2008) using the same routine as with CTx. The WTx measurements were performed after experimenting with different incubation periods, blocker and agonist concentrations. WTx was finally applied only for five minutes, to leave more δ sites unaffected. Additionally the ACh concentration in the pipette was 10 µM to increase the number of events.

### Low-noise modifications

An Axopatch 200B in conjunction with a capacitive feedback headstage (Molecular Devices) was connected to patch clamp pipettes to record single channel currents. Because of its superior electrical and surface characteristics, thick walled quartz glass (outer diameter 2 mm, inner diameter 1 mm) was used to produce pipettes (Parzefall *et al*., 1998; Dudel *et al*., 2000). With modifications described in Parzefall *et al*. (1998), a very low noise level was achieved: 1.5 ± 0.1 (SD) pA root mean square (RMS) at 60 kHz (−3 dB) low pass filtering and 153 ± 30 (SD) fA RMS at 5 kHz (−3 dB) low pass filtering.

### Data storage and filtering

Currents were digitised with a Power 1401 mkII (CED) analogue-digital converter and stored as described in Stock *et al*., (2014). Further processing and analysis were performed with the DC analysis software suite (David Colquhoun, University College London). Before idealisation, recordings were digitally low-pass filtered at 40 kHz, -3 dB cut off, which results in a rise time T_10-90_ = 8.5 µs. Then, recordings were down-sampled from 1 MHz to 500 kHz in Filtsamp. Thus, the sampling frequency was 12.5 times higher than the -3 dB low-pass filter cut off, e.i. 12.5-fold oversampling.

### Data idealisation

To gain high quality dwell time distributions, recordings were idealised using the time course fitting procedure of the Scan program (DC software) (Colquhoun & Sigworth, 1995) until 15,000 transitions were reached. The time course fitting procedure allows to reliably detect events below the rise time of the filter, which is 8.5 µs. The trigger for event detection was set to 3 µs. A final resolution of 5 µs was applied to idealised data (see Results, section High resolution recordings). Due to low open probability and high abundance of single openings, some measurements at low concentrations (10 nM, 100 nM) or measurements after blockade of one of the binding sites did not reach 15,000 transitions. Recordings with unstable current amplitudes were not used for further analysis.

### Dwell time distributions

Dwell time distributions were calculated from idealised traces using Ekdist (DC software). The abscissae were logarithmically transformed to cover the wide range of dwell times. Exponential probability density functions (pdf) were fitted to dwell time distributions and since the ordinate was square-root transformed, peaks in the resulting plot represent time constants of the pdf (Sigworth & Sine, 1987; McManus *et al*., 1987, Colquhoun & Sigworth, 1995). Further details are given in Hallermann *et al*., 2005 and Stock *et al*., 2014.

### Statistics

If not stated otherwise, mean values are reported as mean ± SEM (standard error of the mean). If more appropriate, values are reported as mean ± SD (standard deviation). To choose the number of pdf components for open period, single opening and burst length distributions, the number of components was increased until the probability of erroneously accepting a non-existing component was below 1 % (Rao, 1973; Horn, 1987; Hallermann *et al*., 2005)

## RESULTS

### High resolution recordings

Currents through single muscle type adult nAChRs had a fixed amplitude of -15.0 ± 0.4 pA (Fig. 1), The signal-amplitude allowed low-pass filtering at 40 kHz, while the signal to noise ratio remained at least 11.8 and mean signal to noise ratio was 14.2 ± 1.6. Hence, temporal resolution could safely be set at 5 µs.

**FIGURE 1.**
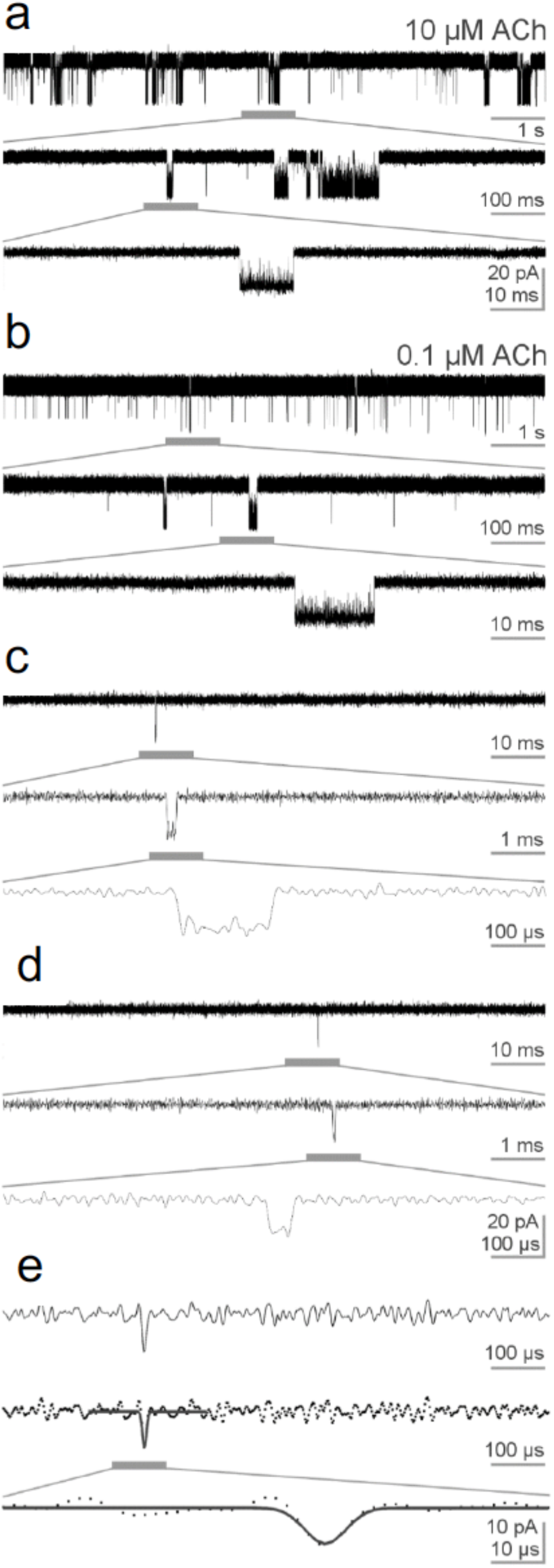
Examples of acetylcholine (ACh) elicited channel openings at high temporal resolution. (*a-e*) Single channel currents recorded at 22°C with thick walled quartz pipettes in the on-cell patch-clamp configuration from transiently transfected HEK 293 cells. The traces are low-pass filtered at 40 kHz and acquired with 10 µM ACh (*a*) or 0.1 µM (*b-e*). Sections of interest are underlined with horizontal grey bars and displayed below with tenfold expanded time scale. Fig,1 *a* and *b* show typical long bursts with long openings, *c* an intermediate 170 µs duration opening, *d* a 45 µs short opening, and *e* a very short 5 µs opening with a time course fit idealisation shown as a solid black line (Scan DC Software).

Fig. 1 shows currents at 40 kHz. Specific events (openings are downward) were marked with a grey bar and shown with a ten-fold expanded time scale below. The 10-second-long upper trace in *a* presents dense groups of channel openings elicited by 10 µM ACh. The marked group (shown expanded below) contains one or two clusters (Colqhoun & Ogden 1988) of channel openings. A typical long or bi-liganded burst of openings is shown below. At 0.1 µM ACh, sections *b-e*, channel opening frequency was much lower than at 10 µM. One of the events in *b* turns out to be again a bi-liganded burst, similar to the one in *a*. The signal in *c* is a single opening of 170 µs, and the highlighted signal in *d*, a single opening of 45 µs duration. Fig. 1 *e* demonstrates the shortest openings at the limit of our resolution. The calibration of the amplitudes was raised to 10 pA and time calibration to 100 or 10 µs for this panel. The short opening is idealised in *e*_*2*_ by the time-course fit procedure of Scan (DC Software) and registered as a 5 µs opening (solid line) which is expanded below. The openings in *c-e* represent examples of three classes of openings. In the following we refer to different types of openings as (τ_o1_) or very short openings, (τ_o2_) short, (τ_o3_) intermediate and (τ_o4_) long openings. Long openings (τ_o4_) are characteristic for bi-liganded bursts of adult nAChRs.

The 5 µs resolution is at its limits in case of very short open or shut times. Fig. 2 shows channel closings (µB) during a bi-liganded burst with the fit in Scan. The last of the three closings shown is 6 µs long, above the temporal resolution, reaches about 10 pA, which is 2/3 of the full amplitude of long events. This shut time and also those down to 5 µs will be represented as results in Figs. 3 to 7. Fig. 2 further shows a 3 and a 4 µs long closing that precede the 6 µs event. Relative to the 6 µs closing the amplitude of these closings are further reduced, and statistics predict that not all events with such durations are resolved. Therefore, below the resolution limit of 5 µs we have to be content with extrapolations.

**FIGURE 2.**
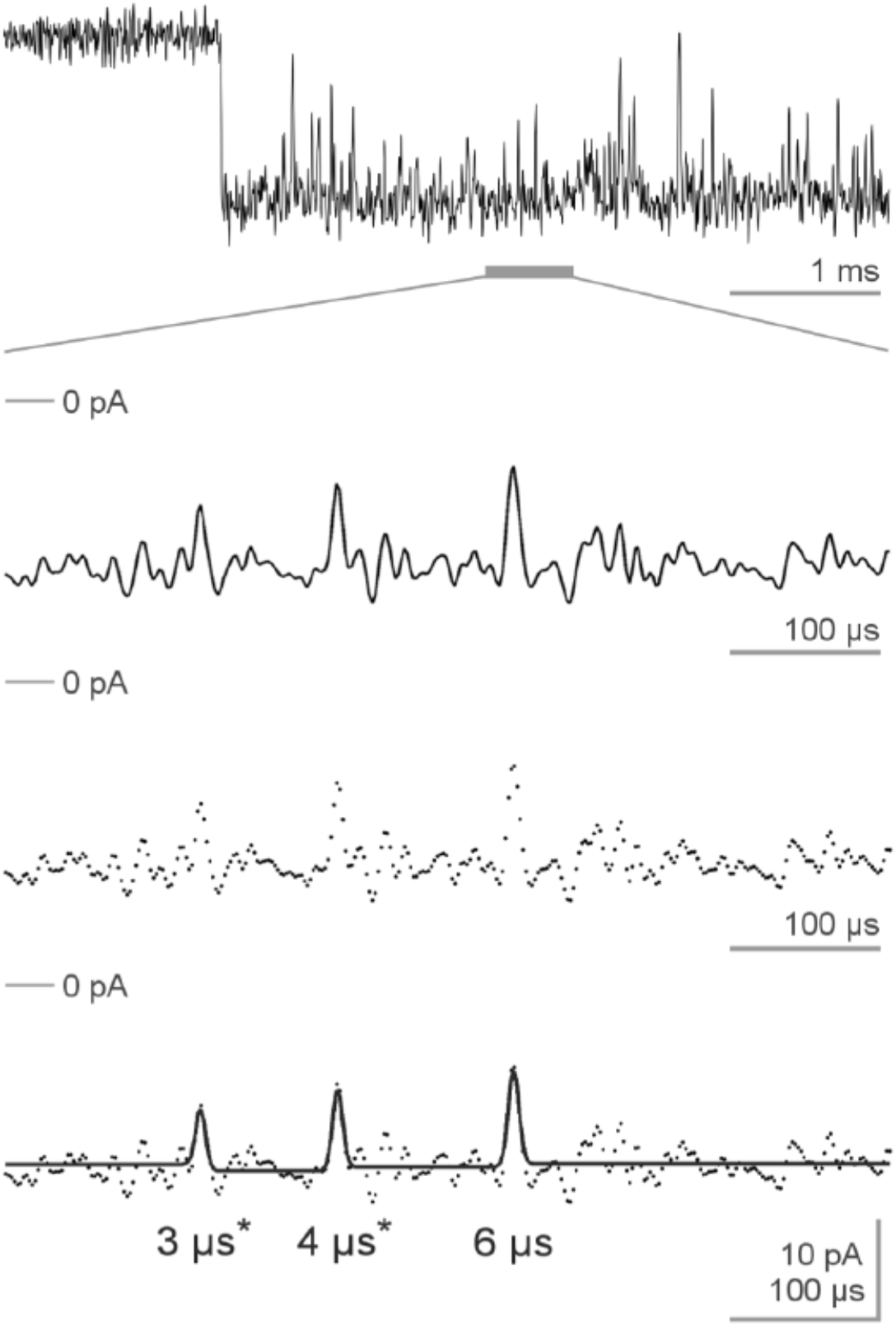
Examples of channel closings during a long burst elicited by 0.1 µM ACh. The tenfold expanded section happens to contain 3 gaps. Above the lower traces, zero current levels are indicated. The traces below illustrate the time course fit idealisation like in Fig. 1 E.

**FIGURE 3.**
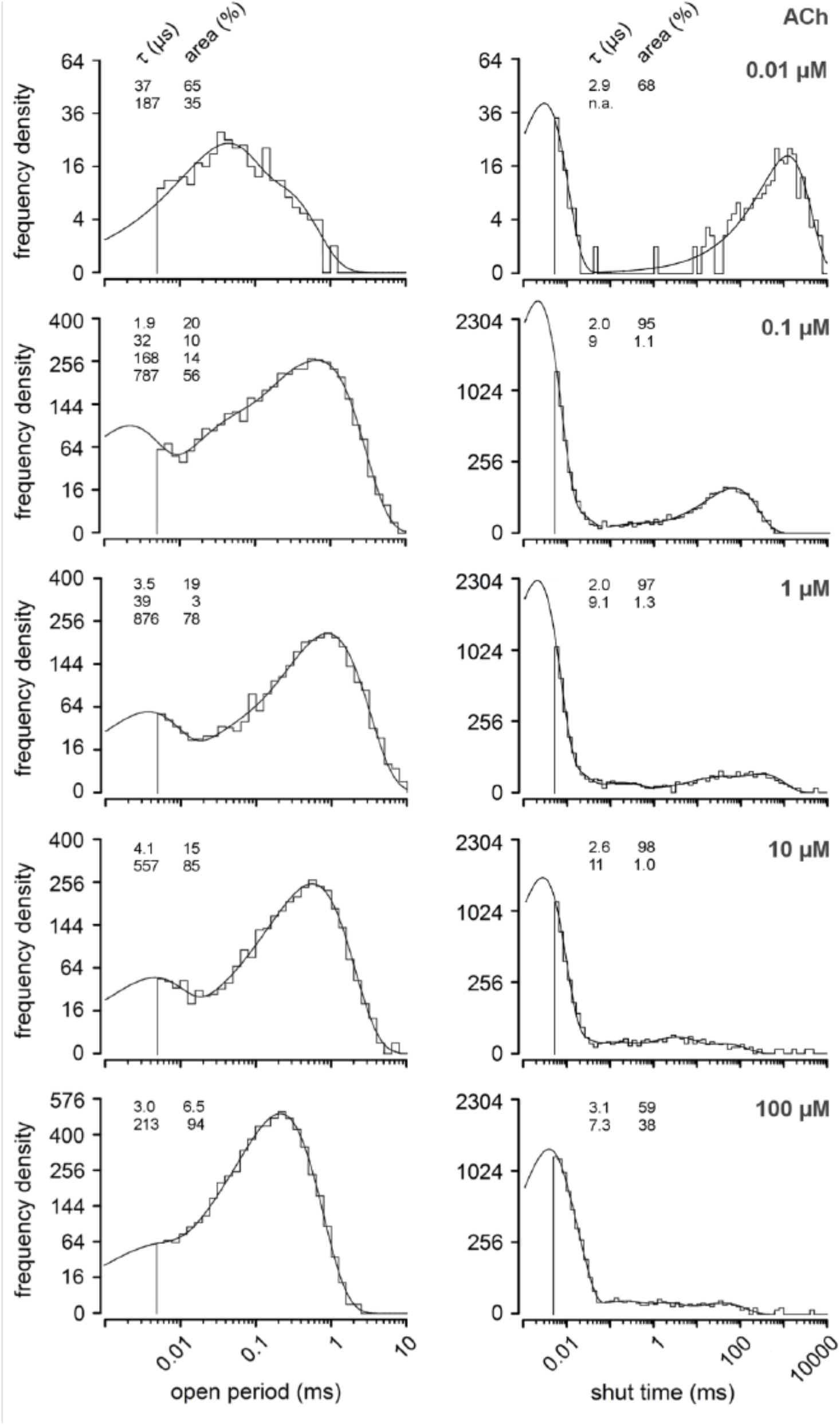
Probability density functions of channel openings and closings with 0.01 to 100 µM ACh. Open periods are plotted on the left and shut time distributions on the right. ACh concentration increases from top to bottom as indicated on the right. Ordinates give number of events per bin and the axis of abscissae event durations on a log scale. Distributions of open periods were fitted by 2 - 4 component probability density functions (see Methods). The parameters τ_o_ and areas of each fit are given in the upper left of each plot. Very short and long openings appear above 0.01 µM ACh concentration. Corresponding distributions of shut times are shown on the right. Each shut time distribution contains a prominent first peak at about 2 µs.

### Open periods and shut times with acetylcholine and epibatidine

Fig. 3 shows distributions of open periods and shut times from 5 sample experiments for a 10 000-fold range of ACh concentration. At 0.01 µM ACh the open periods contain only 2 components (τ_o2_ and τ_o3_), peaking at 37 and 187 µs. The shut times present a large peak at 2.9 µs that reminds us of the breaks seen in the bi-liganded bursts of Fig. 1 *a* *and b*. The second peak at about 1 s reveals that openings are very rare at this extremely low agonist concentration. At 0.1 µM ACh two additional opening components (τ_o1_ and τ_o4_) appear at 1.9 and 787 µs. The first component contains openings like the very short opening in Fig. 1 *e* (τ_o1_). Examples of long openings (τ_o4_) are shown in the bi-liganded burst of Fig. 1 *a*. At 1 µM ACh short openings (τ_o2_) are less frequent and intermediate (τ_o3_) open periods of about 200 µs are not resolved as a separate component. Some openings of this duration may have been obscured by the very large peak of long openings (τ_o4_) at 876 µs and thus not have reached the level of statistical significance.

At higher ACh concentrations (10 and 100 µM) the picture changes further. Again, only two components of open periods are resolved. The very short openings (τ_o1_) are diminished in frequency, and long openings (τ_o4_) shortened: from 876 µs at 1 µM ACh to 557 µs at 10 µM ACh and to 213 µs at 100 µM ACh. At the latter concentration the dominating 2 to 3 µs shut time is joined by a companion, a 7 µs component that is almost as prevalent as the briefest component (see Discussion).

Similar to Fig. 3, open periods and shut times for the agonist epibatidine (Ebd; Prince and Sine, 1998) are shown in Fig. 4. Ebd seems to have a higher affinity to the receptor than ACh, since other than ACh at 0.01 µM it does produce a significant number of long openings (τ_o4_), which stem from bi-liganded nAChRs.

**FIGURE 4.**
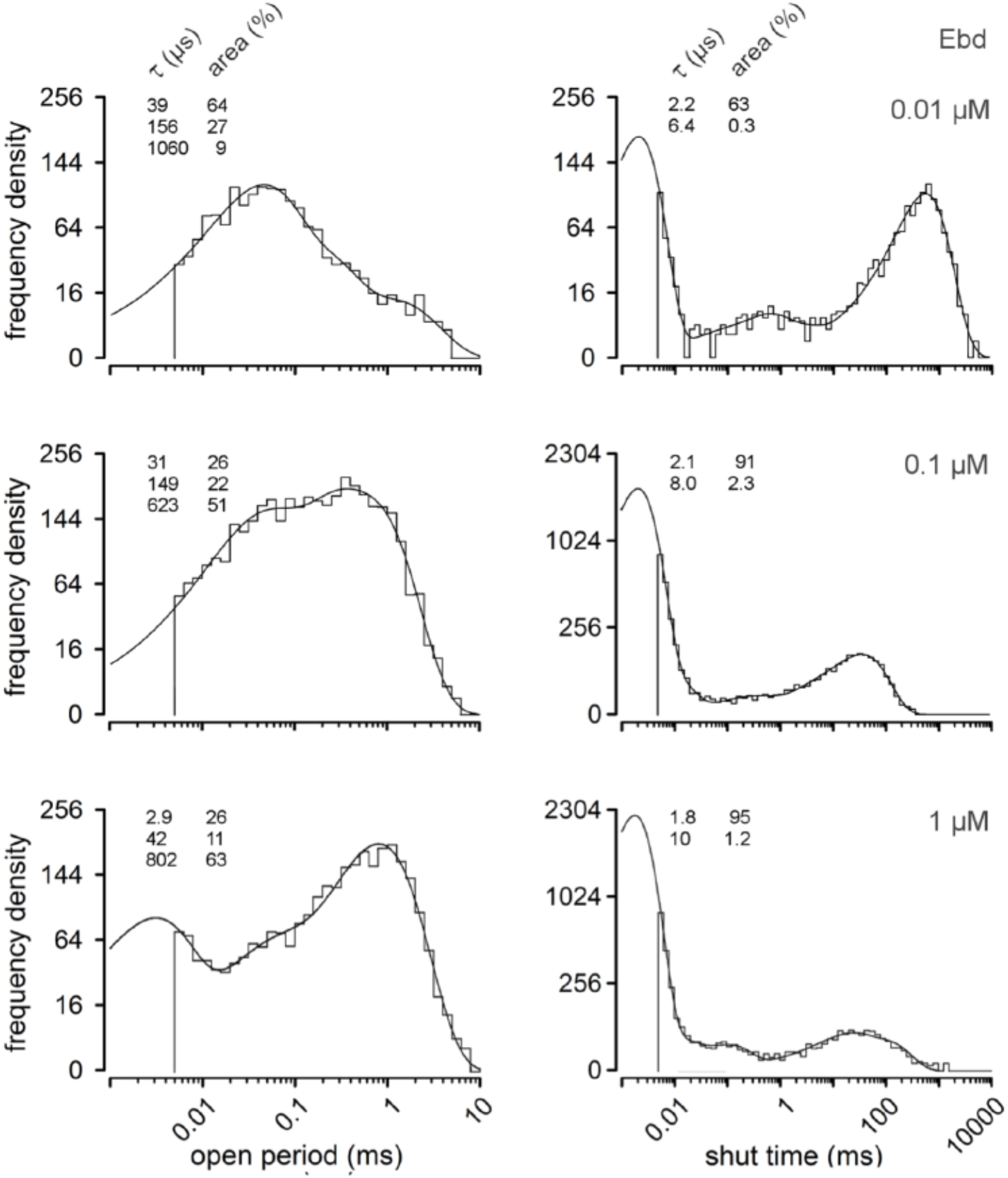
Probability density functions of channel openings and closings elicited by 0.01 to 1 µM epibatidine (Ebd). Same arrangement as in Fig. 3. While long openings appear already with 0.01 µM Ebd very short openings require higher Ebd concentrations. Shut time distributions contain again a prominent first peak at about 2 µs.

The components of 39 and 150 µs are equivalents of τ_o2_ and τ_o3_ seen with ACh. At all concentrations of Ebd in Fig. 4, a very short shut time is prominent. With 0.1 µM Ebd again 3 types of openings appear, but now the long component (τ_o4_) dominates. Different from the same ACh concentration, the very short open period (τ_o1_) is not resolved. At 1 µM Ebd, very short open periods are present (like with ACh), but again like with ACh, τ_o3_ open periods are not resolved.

### Mono- and bi-liganded bursts of openings

Openings of nAChRs appear either as single openings separated by long shut times or as bursts i.e. openings separated by short closings (Colquhoun & Sakmann 1981). A critical shut time (t_crit_) is used to distinguish between bursts and single openings. All openings separated by closings shorter or equal t_crit_ are considered as openings within one burst. A closing longer than t_crit_ ends a burst. Openings flanked from both sides by closings longer than t_crit_ are considered single openings. The duration of t_crit_ was obtained by extrapolating the first shut time component and set to 26 µs with ACh and 20 µs with Ebd. The most prominent and best studied activity in single channel recordings of muscle type nAChRs are long bursts, generally they are thought to arise from nAChRs with two agonists bound. They consist of long openings (τ_o4_) that are separated by very short 2 µs shut times. There are also short bursts that appear for instance in the recording for Fig. 3 at 0.01 µM ACh (illustrated in some detail in Fig. 6). At this concentration only mono-liganded openings are elicited, as ascertained by the absence of bi-liganded long bursts. Mono-liganded bursts consist of τ_o2_ and τ_o3_ (or rarely τ_o1_) openings separated again by very short 2 µs shut times. Plots of the durations of mono- and bi-liganded bursts versus ACh concentrations are shown in Fig 5. The duration of the bi-liganded bursts decreases with increasing concentration for both agonists (see Discussion).

**FIGURE 5.**
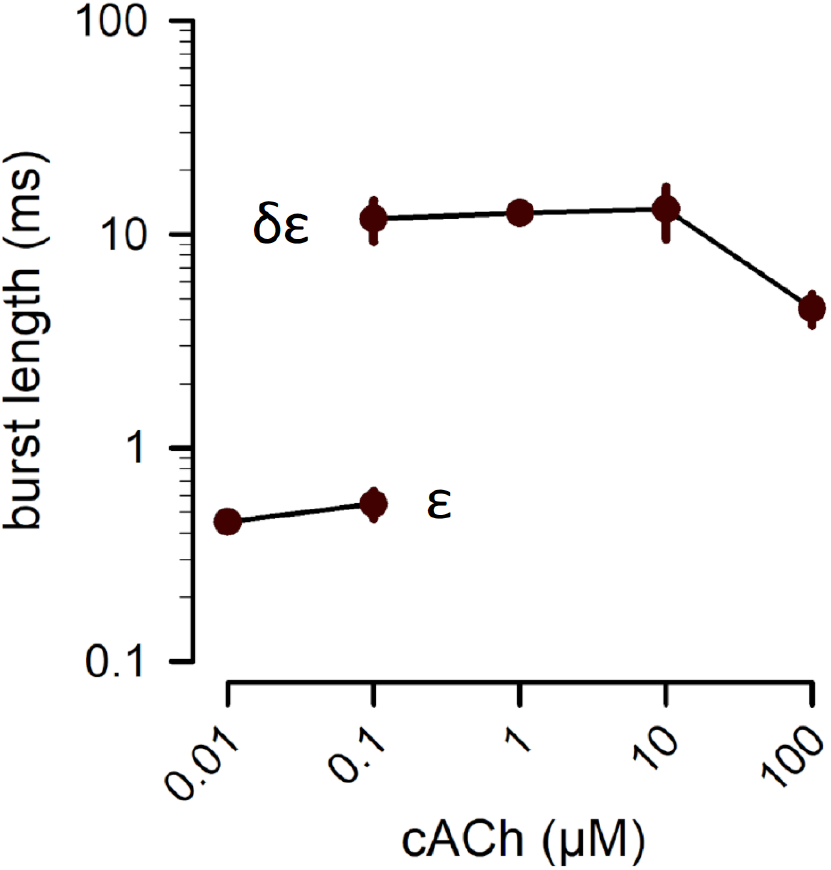
Concentration dependence of the duration of mono-liganded (ε) and bi-liganded (δε) bursts. Mean duration ± SEM in recordings with ACh (3 recordings for each concentration).

**FIGURE 6.**
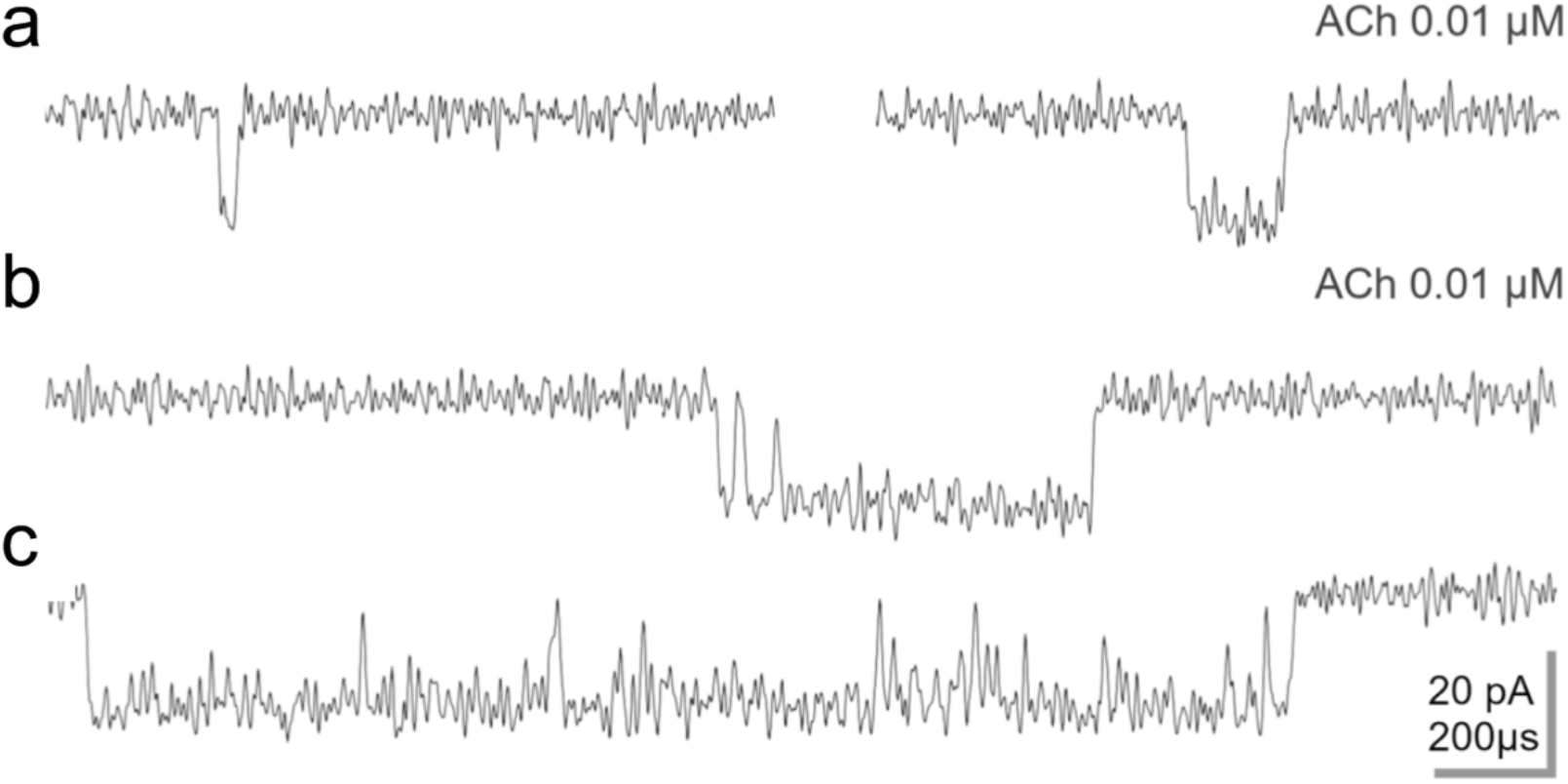
Mono-liganded bursts elicited by ACh. (*a*) Examples of mono-liganded single openings with 0.01µM Ach, low-pass filtered at 60 kHz. (a)The short event on the left is 31 µs and the intermediate one on the right 107 µs long. (*b and c*) Examples of two mono-liganded bursts with 0.01µM ACh again low-pass filtered at 60 kHz.

Fig. 6 presents examples of various mono-liganded bursts of channel openings at low agonist concentrations. Fig. 6*a* shows a short (τ_o2_) and an intermediate opening (τ_o3_) from an experiment filtered at 60 kHz. Interestingly, short and intermediate openings both occur in what may be termed mixed mono-liganded bursts. An example of such a mono-liganded burst with two short openings at the start, followed by an intermediate opening is illustrated in Fig. 6*b*. A relatively long mixed mono-liganded burst with a succession of intermediate openings followed by short openings is shown in Fig. 6*c*.

### Blocking the δ- or the ε-site of nAChRs

We blocked the δ-site by incubating cells in 1 µM of the snail *α-Conotoxin M1* (CTx, see methods and Fig. 7 *a,b*). This should reveal channel activity generated by agonist binding at the ε-site of nAChRs. Both ACh and Ebd elicited short single openings (τ_o2_) but no intermediate openings (τ_o3_) and 120 - 130 µs mono-liganded bursts of openings (Fig. 7 *a,b*). Mono-liganded bursts with CTx are somewhat shorter than mono-liganded bursts without blocker and contain only short (τ_o2_) openings (Fig. 6 *a,b*). Shut times within bursts show again a prominent very short < 3 µs component.

**FIGURE 7.**
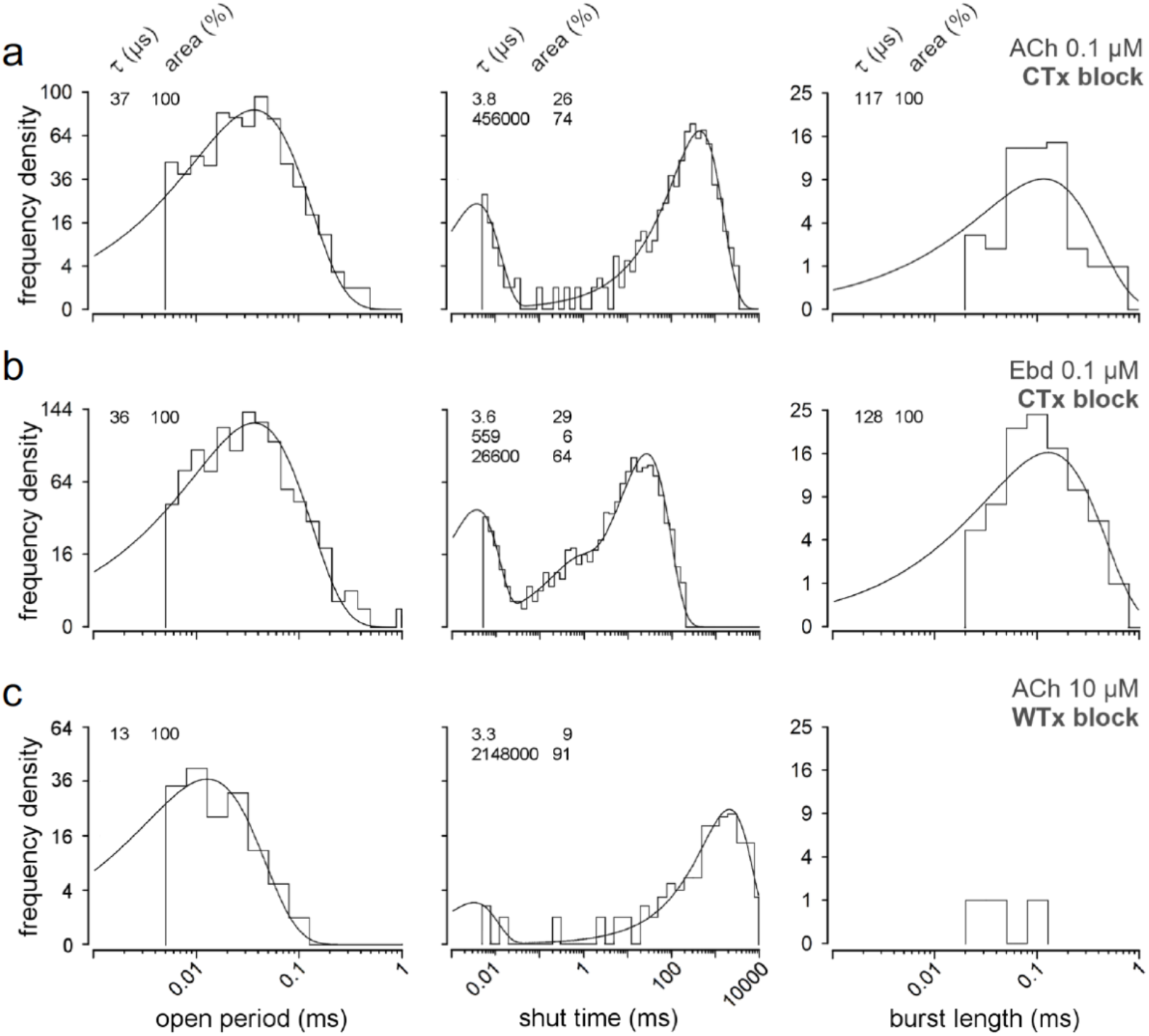
Dwell time distributions from ACh or Ebd recordings after incubation in *α-Conotoxin-M1* (CTx) *or* Waglerin-1 (WTx), specific blockers of the δ-site and ε-site, respectively. Open period, shut time and burst length distributions with 0.1 µM ACh (*a*) or Ebd (*b*), after incubation with 1 µM CTx. Open period distributions were fitted with a one component pdf. Shut time distributions were fitted with either two (ACh) or three (Ebd) components. Burst length distributions were fitted with one component. Time constants of the fits (τ; µs) and their proportion (area; %), are shown on the upper left of each plot. (*c*) Open period, shut time and burst length distributions with 10 µM ACh after incubation with 1 µM WTx.

Waglerin-1 (WTx) is a snake venom that blocks neuromuscular transmission in mammals (Sellin *et al*., 1996; McArdle et al., 1999; Molles *et al*., 2002) preferentially at the ε-site of nAChRs. Data for a cell incubated with 1 µM WTx are shown in Fig. 7c. The shut time-plot indicates that openings are extremely rare, on average less than one occurred during one second. Furthermore, only very short openings occurred with a mean duration of 13 µs. Almost all openings were singles but occasionally bursts occurred as illustrated in the right panel of Fig. 7c.

### A net of states

Fig. 8 shows a representation of the reactions of the receptor (R) with agonists (A). Binding of A to site Rε starts a burst of repetitive channel openings µB – Oε. Binding to the other site (Rδ) or both sites simultaneously (Rδε) would elicit analogous reactions.

**FIGURE 8.**
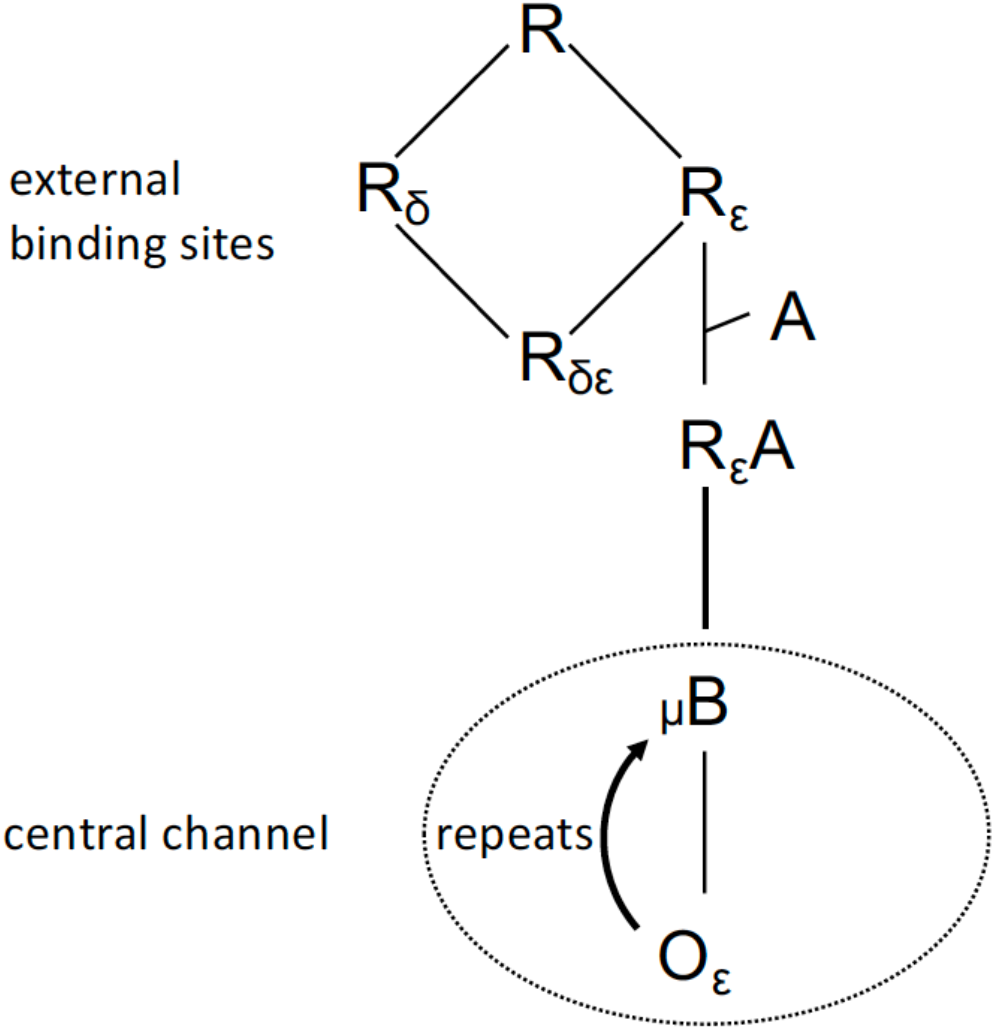
Network of states of the AChRs molecule exposed to the agonist ACh. The receptor (R) binds the agonist (A) to its binding site (Rε) to form (RεA) that connects to the central channel of the molecule initiating openings (Oε) that are bordered by short channel blocks (µB).

## DISCUSSION

### Functions of binding sites

The first aim of the present study was to identify the functional roles of the transmitter binding sites of nAChRs. In the ring of subunits agonists can bind at the interfaces between α- and δ- as well as α- and ε -subunits in short at Rδ and or Rε (Fig. 8). Simultaneous activation of δ- and ε-sites (to OA_2_) elicits bi-liganded bursts of τ_o4_ openings (Colquhoun & Sakmann 1985). We were especially interested in the roles of the mono-liganded activations to gain Oδ or Oε. The agonists ACh and Ebd have high affinity for the Rε-site. When ACh is applied at the very low concentration of 0.01µM, only τ_o2_ and τ_o3_ appear (Fig. 3), while with 0.1 µM ACh all four types of openings are generated. Thus, τ_o2_ and τ_o3_ are elicited at ε-sites, and τ_o1_ at δ-sites. This conclusion is supported by the mixed τ_o2_ and τ_o3_ bursts in Fig. 6, showing instantaneous switches between τ_o2_ and τ_o3_ openings, indicating that τ_o2_ and τ_o3_ openings are generated from the same binding-site. Blocking the δ-site by CTx resulted in τ_o2_ openings and blocking the ε-site by WTx led to very short openings that may resemble τ_o1._

### Functions of the center of the molecule

The center of the nAChRs molecule contains a channel that may pass ionic current across the cell membrane (see Introduction). Astonishingly, the amplitude of the open channel current is strictly fixed to 15 pA. In addition, the current flow is interrupted by strictly fixed 2-3 µs short full closings (µB) that border the openings of the channel (Fig. 8). These reactions of the channel are elicited by a signal from the binding site that has been activated by binding the agonist. This signal also determines the duration of the specific channel opening Oε (Fig. 8).

All these functions of the nAChR molecule are united to form bursts of channel openings. The description of the functions of the nAChR molecule seems to be valid also for different agonists (Suberyldicholine, Epibatidine, Carbachol) at the adult as well as the embryonic nAChRs (Parzefall et al. 1998; Hallermann et al., 2005; Stock et al. 2014).

### Limited time resolution of records

In our recordings we can only resolve currents down to 5 µs duration and extrapolate dwell times down to < 3 µs. The shortest shut times should be much shorter than measured presently (Fig. 2), and the yet unresolved ones would frequently break up the measured openings into very short open times and intermittent closings. Probably the “< 3 µs” µB shut times are generated by some rhythmic movement of electrical charges in the wall of the channel. Such movements would continue as long as the channel is open due to binding of an agonist at the δ- or ε -site, and thus would transform all the channel currents into continuous high frequency bursts of openings, each started by a channel closing (µB) and finished by a channel closing (µB). With a very special technique and high bandwidth Hartel et al. (2018) demonstrated continuous bursts with about 1 MHz frequency from an intracellular Ca^2+^-channel. Molecular dynamics simulations of neuronal α4β2 nAChRs show movements below 1 µs (Oliveira et al., 2019). These techniques may be extended to adult muscle type nAChRs and could eventually perhaps be related to the types of channel openings and < 3 µs µB shut times described here.

### Desensitization

The present recordings were generated by single nAChR molecules that react to ACh or Ebd, as depicted in Fig 8. Patch-clamp experiments with fast application of agonist on nAChR molecules result in sums of currents from hundreds of such molecules (Franke et al., 1991). When the patch is exposed to a pulse of ACh at a concentration > 1 µM, the elicited initial current pulse decays exponentially to a lower maintained level, it is “desensitized”. An ACh concentration of 100 µM will desensitize the current from the patch by a factor of about 100. In our experiments on single receptor molecules we expose the receptor continuously to 100 µM ACh, but we see no signs of desensitization: The recorded current amplitude measures the standard 15 pA. Obviously, desensitization reduced the number of activatable receptors.

At the highest concentration of 100 µM ACh that was applied in this study, the dominating 3 µs shut time was joined by a relatively large 7 µs component (Fig. 3). This component tended to increase at higher ACh concentrations. A concentration-dependent component should be initiated by the binding of ACh, but we have no information about its position in the network of Fig. 8. We mention this unclear finding, since it may relate to open channel block at high ACh concentrations.

## CONCLUSIONS

With the same techniques as the present ones, also the juvenile type of nAChRs has been investigated (Stock et al. 2014). The results were very similar to those for the adult nAChRs, only the current amplitudes were lower than 15 pA and the discussion was less detailed. The main function of the nAChRs is transmission of the impulse of the motor nerve to depolarization and excitation of the muscle fiber by long bursts of openings mediated through the bi-liganded receptor Rδε. The functions of the mono-liganded receptors are unclear. At very low ambient ACh concentrations of the serum the relatively small currents generated by mono-liganded bursts of openings might help to establish and stabilize the nerve-muscle synapse (Xie and Poo, 1986; Bloch-Gallego, 2015).

## ABBREVIATIONS (defined in the text at first mention)

ACh: acetylcholine
δ-site: αδ-ligand binding site of muscle-type nAChRs
ε-site: αε-ligand binding site of muscle-type nAChRs
ε-burst: short burst
εδ-burst: long burst
µB: block of central channel for < 3 µs
CMOS: complementary-metal-oxide-semiconductor
CTx: α-Conotoxin-M1
Ebd: Epibatidine
EPSC: excitatory postsynaptic current
nAChR: nicotinic acetylcholine receptor
n_corr_: corrected number of short gaps per burst
pdf: probability density function
RMS: root mean square
SEM: standard error of the mean
SD: standard deviation
t_crit_: critical shut time
τ_o_: time constant tau of open period probability density function
WTx: Waglerin-1

## AUTOR CONTRIBUTIONS

D.L., J.D., M.H., designed the experiments. D.L. performed the experiments. D.L., A.M., M.P., J.D. and M.H. analysed the experiments, contributed to writing and revising the paper. All authors approved the final manuscript for publication and agree to be accountable for all aspects of the work in ensuring that questions related to the accuracy or integrity of any part of the work are appropriately investigated and resolved. All persons designated as authors qualify for authorship, and those who qualify for authorship are listed.

## ACKNOWLEDGEMENTS

This work was supported by grants from the Deutsche Forschungsgemeinschaft (DFG, German Research Foundation) TRR 166 Receptor Light B06, FOR 3004 SYNABS P1.

